# Strong promoters are mutationally robust

**DOI:** 10.1101/2025.10.20.683477

**Authors:** Timothy Fuqua, Stepan Denisov, Mato Lagator, Andreas Wagner

**Author notes:** Authors contributed equally (Stepan Denisov, Mato Lagator).

## Abstract

Mutational robustness is the persistence of a phenotype upon mutation. It facilitates molecular evolution and has been characterized in a variety of biological systems, but studies of prokaryotic promoters are limited. Prokaryotic promoters are non-coding DNA sequences that regulate transcription. The main housekeeping promoters (σ70) share two sequence motifs called the -35 and the -10 box that are spaced 17±1 base pairs (bps) apart. The sequence of these boxes, their distance, and the existence of multiple nearby boxes can determine promoter strength, the ability of a promoter to drive high levels of transcription. Here we used computational modelling to show that the mutational robustness of σ70 promoters – the persistence of promoter strength upon mutation – correlates with promoter strength itself. Mutational robustness is also influenced by potential overlaps between -10 and -35 boxes, and by specific positions within the boxes. It is higher when the boxes are exactly 17 bps apart. These findings can be partially explained by the flexibility of -35 boxes, which enables adjacent bases to create overlapping and new -35 boxes. Our work can help to engineer synthetic promoters for strength and robustness, and to understand the dynamics of promoter evolution.

## Introduction

Bacterial promoters are DNA sequences that serve as docking sites for RNA polymerase to initiate transcription (Huerta and Collado-Vides 2003; Paget 2015). The textbook example of the housekeeping promoter (σ70) encodes two motifs for RNA polymerase called the -35 box (consensus TTGACA) and the -10 box (consensus TATAAT), spaced 17±1 bp apart (Pribnow 1975; Deal et al. 2024). Many promoters deviate from this consensus. For example, spacers have been described outside of this 17±1 bp range (Wang et al. 2020; Klein et al. 2021), and there is substantial variation in -10/-35 box sequences (Urtecho et al. 2019; Wang et al. 2020). This diversity influences promoter strength (Lagator et al. 2022; Fuqua et al. 2024; Fuqua and Wagner 2025), but it is unclear how it relates to mutational robustness.

Mutational robustness is the extent to which a phenotype persists upon mutation (Rennell et al. 1991; Montville et al. 2005; Wagner 2007). It can counterintuitively stimulate evolution. This is because a more robust phenotype can accumulate more genetic variation via drift (Schuster et al. 1997), allowing an evolving population to explore a greater proportion of sequence space, thus providing greater access to more novel phenotypes (van Nimwegen et al. 1999; Wagner 2007; Wagner 2012). Selection can also directly act upon mutational robustness (Montville et al. 2005; Wagner 2005; Fares 2015), particularly in species with large effective population sizes (N_e_) and/or high mutation rates (µ) (Draghi et al. 2010).

The ability of a promoter to maintain gene expression upon mutation, i.e. mutational robustness, is important for evolution (Einarsson et al. 2022). Studies which explore the mutational robustness of promoters have found that transcription factors can buffer the effects of mutations in core promoter sequences (Parisutham et al. 2025). Another recent study demonstrates that core promoter sequences can differ in their mutational robustness (Tsuru et al. 2024), but did not explore the sequence features of such robustness.

Here, we use computational modeling to show that mutational robustness directly correlates with promoter strength. Normalizing mutational robustness to promoter strength also reveals differences in robustness independent of promoter strength. For example, promoters with exact 17 bp spacers, overlapping -10 and -35 boxes, and specific -10 and -35 box sequences are more mutationally robust than other promoters. The reason for promoter mutational robustness may lie in the flexibility of -35 boxes, which allows adjacent bases and mutations to encode additional -35 boxes 1 bp upstream or downstream of them. Because of the intrinsic relationship between promoter strength and robustness, we also argue in the Discussion that promoter robustness may be a selected trait.

## Results

### Mutational robustness positively correlates with predicted promoter strength

Position weight matrices (PWMs) are computational tools used to calculate a score in information-theoretical units (bits) for how well a query sequence matches a set of experimentally derived binding sites for a specific protein, such as RNA polymerase. They are used to identify putative binding sites if the query sequence score is above a given threshold (Berg and von Hippel 1987; Hertz and Stormo 1999). PWM scores for -10 and -35 boxes can be used as a proxy for promoter strength – how much mRNA is transcribed from a promoter - as their scores correlate with experimentally validated promoter sequences (**Fig S1A**, see also (Bharanikumar et al. 2018; Fuqua and Wagner 2025)).

We asked if PWMs could also be used to quantify the mutational robustness of promoters. To this end, we computationally generated a set of all possible 6-mer DNA sequences (-10 and -35 boxes are 6 bp), and identified which of these 4,096 sequences qualify as putative -10 and -35 boxes (Tierrafría et al. 2022) based on how their scores relate to their respective PWM’s information content (Patser threshold, see Methods). We identified 39 sequence instances that classify as a -10 box (**Fig 1A**). For each instance, we calculated the percentage of single mutational neighbors that are also instances (18 neighbors for each instance). In other words, we calculated how likely it is that a -10 box remains a -10 box after a single point mutation. We refer to this quantity as the mutational robustness of a -10 box.

**Figure 1.**
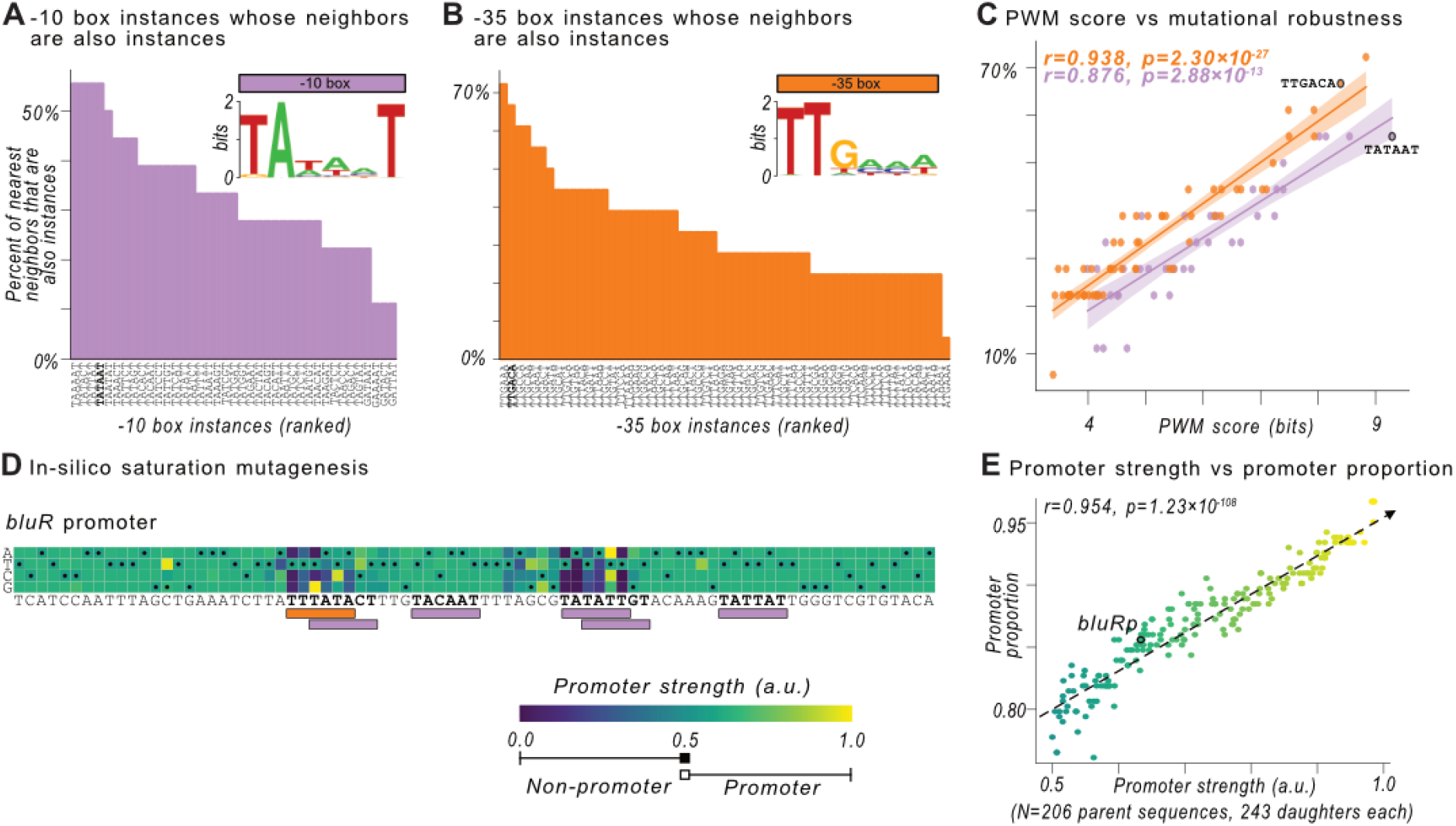
Mutational robustness positively correlates with predicted promoter activity. **(A)** Top right: a sequence logo generated from a position-weight matrix (PWM) for the -10 box. Bar plot: we calculated all possible DNA sequence instances that the PWM classifies as a -10 box (see **Methods**). For each instance of such a box, we plot the percentage of its nearest mutational neighbors (out of 6×3=18 possible neighbors) that are also -10 box instances. **(B)** Analogous to (A) except for the -35 box. In both panels (A) and (B) consensus instances of the respective boxes are shown in bold on the x-axis. **(C)** The PWM scores in bits for each box instance vs the percentage of a box’s nearest neighbors that are also an instance. Orange circles: -35 box instances, magenta circles: -10 box instances. We tested the null hypothesis that there is no correlation between the percentages and PWM scores using a Pearson correlation (-35 box: r=0.938, p=2.30×10^-27^; -10 box: r=0.876, p=2.88×10^-13^). **(D)** We created in-silico saturation mutagenesis libraries (a library of all possible single-mutants from a given sequence) for 206 promoter sequences of length 81 bp (i.e., 243 = 81 × 3 single point mutant sequences for each promoter sequence). For each single point mutant, we predict the promoter score in arbitrary units (a.u.) using a thermodynamic model (Lagator et al. 2022). We call sequences with scores less than or equal to 0.5 a.u. “non-promoters” and those greater than this value “promoters.” We exemplify these predictions for the *bluR* promoter and its mutations as a heatmap matrix, where the color of each corresponding matrix entry to the predicted promoter scores for mutations at each position (x-axis) and base (y-axis). The wild-type sequence is written underneath the matrix, and the matrix elements corresponding are inscribed with small black circles. Below the sequence are magenta and orange rectangles, which correspond to PWM predictions for the -10 and -35 box, respectively (see A and B). **(E)** Predicted promoter strength in arbitrary units (a.u.) vs the proportion of single point mutant sequences that are still promoters. The color of a circle corresponds to the predicted promoter strength (see color bar in D). We test the null hypothesis that there is no correlation between the strength and proportion (Pearson correlation, r=0.954, p=1.23×10^-108^).

We used the same approach to identify 58 sequence instances of -35 boxes (**Fig 1B**), and each instance’s mutational robustness – the percentage of neighbors that are also -35 boxes. See **Fig S2** for additional information on the -10 boxes (**Fig S2A**) and the -35 boxes (**Fig S2B**).

For each -10 box instance, the percent of nearest neighbors that are also -10 boxes ranges from ∼11% (2/18) to ∼56% (10/18), with an average of ∼33% for each -10 box instance (**Fig 1A**). For the -35 box instances, these values range from ∼6% (1/18) to ∼72% (13/18), respectively, with an average of ∼34% (**Fig 1B**). There is no significant difference in the distribution of these percentages between -10 box and -35 box instances (two-tailed t-test, t=-0.329, p=0.743) (**Fig S2C**). These calculations suggest that -10 and -35 boxes are similarly robust on average, although robustness can vary substantially among individual instances. Using lower PWM thresholds does not change these conclusions (**Fig S3**).

We found that boxes with high mutational robustness also tend to have high PWM scores. This holds for both -10 boxes (Pearson correlation, r=0.876, p=2.88×10^-13^) and -35 boxes (r=0.938, p=2.30×10^-27^) (**Fig 1C**, see **Fig S1A** comparing PWM scores vs putative promoter sequences). Because PWM scores are a proxy for promoter strength (Berg and von Hippel 1987; Bharanikumar et al. 2018), our findings suggest that upon mutation, stronger promoters are more likely to remain promoters, i.e., still drive transcription. Note that this finding is solely for core promoter sequences and ignores potential interactions with activators or repressions.

Previous studies have demonstrated that PWM scores do not perfectly correlate with promoter strength (Urtecho et al. 2019; LaFleur et al. 2022). Additionally, one shortcoming of PWMs is that they do not account for dinucleotide interactions within a binding site (Siddharthan 2010; Zhao et al. 2012). To support our finding that mutational robustness correlates with promoter strength, we also used a statistical thermodynamic model that captures more precisely the impact of each sequence and, importantly, spacer length variation, on the binding energy of RNA polymerase (Lagator et al. 2022). The model outputs the probability of a query sequence being a promoter, which significantly correlates with both PWM scores and experimentally validated promoter activity (**Fig S1B,C**). We directly translate these probabilities into scores that range from 0 (zero promoter strength, no expression) to 1 (maximal promoter strength).

To quantify the mutational robustness of promoters, we first needed a set of promoters for the model to compare. To this end, we acquired a set of 1,997 promoters (81 bp each) from the Regulon DB database (Tierrafría et al. 2022). For these 1,997 promoters, the thermodynamic model predicts that RNA polymerase binding alone is sufficient for expression (from 0.5 – 1.0 a.u.) in 206 of them. These 206 genomic promoters vary in the amount of evidence supporting their validity. Specifically, 88% (182) were identified computationally without human oversight, 47% (97) were validated using transcription initiation mapping, and 36% (74) were identified by human experts based on their proximity to known open reading frames and transcription start sites (see Methods). Importantly, 100% (206) are predicted to be promoters using the thermodynamic model.

We then created in-silico saturation mutagenesis libraries for each promoter sequence, which comprise all possible single point mutations in a promoter (81 bp × 3 other possible bases = 243 single point mutant sequences per promoter), and calculated their respective promoter scores (**Fig 1D**). Finally, we calculated for each promoter and its respective single point mutants, the proportion of point mutants with promoter scores greater than 0.5 a.u., which we refer to as the *promoter proportion*.

Promoter proportions range from ∼76% to ∼97%, with an average of ∼87%. In other words, in these 81 bp sequences, anywhere from ∼3% to ∼24% of single point mutations will disrupt a given promoter. This also means that some promoters are more robust to mutations than others. In addition, the most robust promoters are also the strongest promoters (Pearson correlation between promoter proportion and strength r=0.954, p=1.23×10^-108^) (**Fig 1E**).

In sum (**Fig 1**), two different computational models based on experimental evidence and computational predictions demonstrate a strong and positive correlation between predicted promoter strength and mutational robustness. Note that based on how it is calculated, our definition of robustness is binary, i.e. a sequence is either a promoter (strength > 0.5 a.u.) or not. See **Fig S4** for alternative definitions of robustness such as the coefficient of variation, which decreases with promoter strength (**Fig S4A**) and the standard deviation, which initially increases with promoter strength, but then decreases (**Fig S4B**).

### Mapping promoter architectures

We next wanted to understand how -10 and -35 boxes in the 206 promoters influence their predicted strength. To this end, we systematically map which PWM-predicted -10 and -35 boxes associate with changes in promoter strength, and tested whether significant differences exist in the central tendencies of promoter strengths for mutants with and without the PWM-predicted box using a Mann-Whitney U test (MWU, see **Methods**). We refer to boxes whose removal by mutation significantly decreases promoter strength by more than 0.1 a.u. as *mapped* boxes. As an example, the promoter for a gene called *bluR* is in our dataset (**Fig 2A**). When point mutations remove a PWM-predicted -35 box from the promoter, the predicted promoter strength decreases by -0.286 a.u. (two-tailed MWU test, U=208, q=2.68×10^-6^) (**Fig 2B**). Similarly, when mutations remove one of the PWM-predicted -10 boxes (TATATT), the predicted promoter strength decreases by -0.526 a.u. (two-tailed MWU test, U=13, q=1.92×10^-7^) (**Fig 2C**).

**Figure 2.**
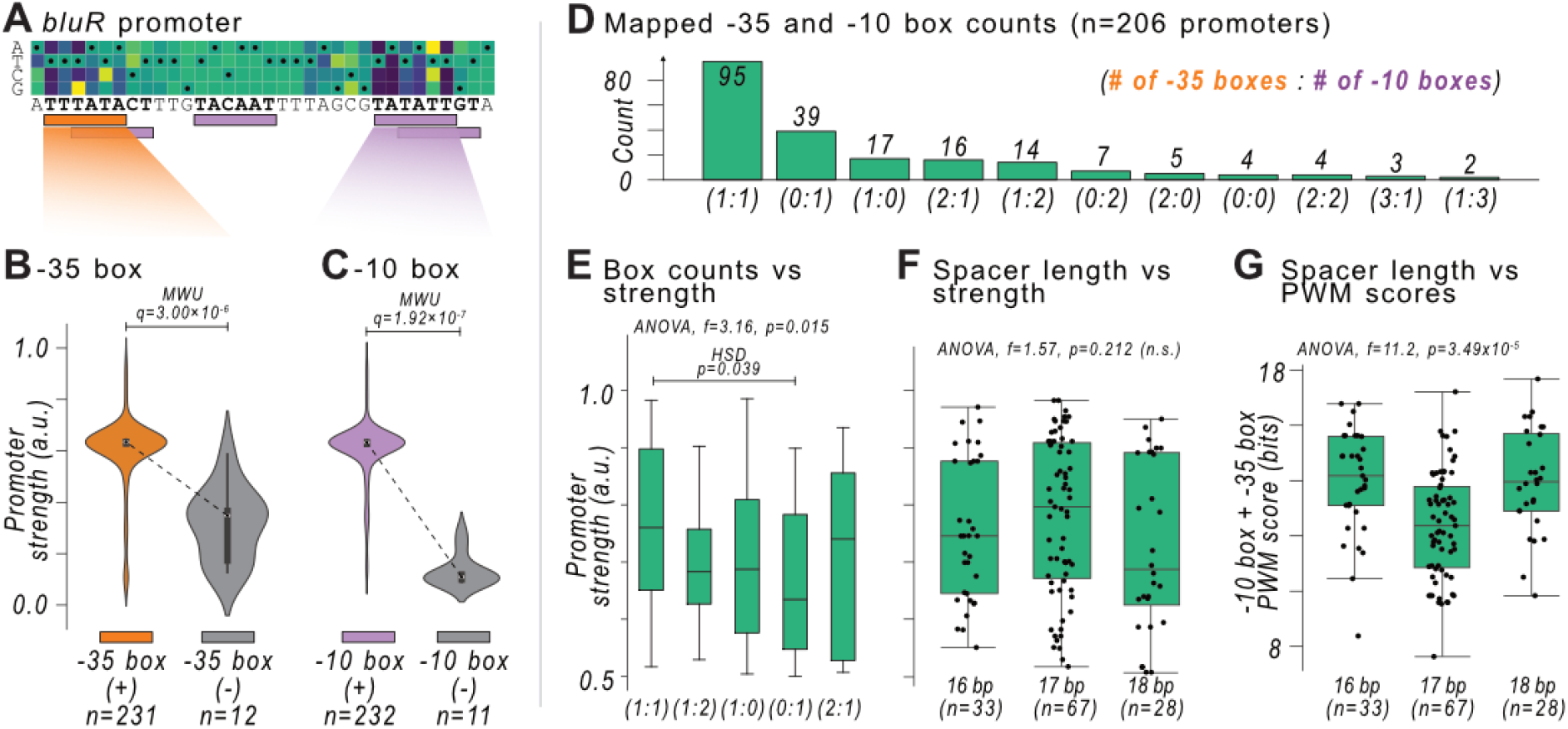
Mapping -10 and -35 boxes reveals diverse promoter architectures. **(A)** Promoter strength predictions from *in-silico* saturation mutagenesis library for the *bluR* promoter. The color of each matrix element corresponds to the predicted promoter strengths for mutations at each position (x-axis) and base (y-axis). The wild-type sequence is written below the matrix and its corresponding matrix elements are inscribed with small black circles. Below the sequence are magenta and orange rectangles, which correspond to PWM predictions for the -10 and -35 box, respectively. **(B)** We test the null hypothesis that mutant sequences with the indicated -35 box (orange, position-weight matrix) have the same promoter strengths as mutant sequences without a -35 box (gray) (two-tailed Mann-Whitney U [MWU] test, q=3.00×10^-6^). Each violin plot corresponds to a kernel density estimate of a strength distribution. **(C)** Analogous to (B) but for the indicated -10 box (two-tailed MWU test, q=1.92×10^-7^). **(D)** We mapped -10 and -35 boxes in 206 promoter sequences as exemplified in (B-C), and categorized promoters by their number of mapped -35 and -10 boxes. Underneath each bar, the left number refers to the number of mapped -35 boxes, and the right number to the number of mapped -10 boxes in the promoters whose number (among 206) is indicated by the bar’s height. **(E)** We tested the null hypothesis that the strengths for promoters in each category are indistinguishable, using an analysis of variance (ANOVA) on architectures with at least 10 promoters each (ANOVA, f=3.16, p=0.015). We used the Tukey HSD post-hoc test (Tukey 1949) to identify a significant difference between category (1:1) and (1:0) (one -35 box and one -10 box vs no -35 box and one -10 box, p=0.039). None of the other pairwise comparisons differed significantly. **(F)** For promoters with a -10 and -35 box spaced 16-18 bp apart (n=128), we used an analysis of variance to test the null hypothesis that the mean promoter strengths for the different spacer lengths are indistinguishable (ANOVA, f=1.57, p=0.212, n.s.= not significant). **(G)** Analogous to (F) but for the sum of PWM scores for the -10 and -35 box in each spacer length category (ANOVA, f=11.2, p=3.49×10^-5^).

We mapped -10 and -35 boxes in the 206 promoters using this approach, finding substantial variation in the number of mapped -10 (209) and -35 (187) boxes. For instance, ∼46% (95/206) of the promoters have both a mapped -10 and a -35 box (**Fig 2D**), ∼19% of promoters have only a mapped -10 box (39/206). The latter percentage is consistent with previous studies which demonstrate that ∼20% of promoters do not encode -35 boxes (Mitchell et al. 2003). Conversely, only ∼8% (17/206) of promoters encode a -35 box but not a -10 box. See **Fig 2D** for data on the remaining 27% of -10/35 box configurations. This analysis demonstrates that promoter architectures are diverse. The canonical architecture (one -10 box and one -35 box) makes up only about half of promoters based on our thresholds and modeling parameters.

We asked whether the number of mapped -10 and -35 boxes per promoter correlates with promoter strength (ANOVA, f=3.16, p=0.015), and found that promoters with a -35 and a -10 box are ∼ 0.1 a.u. stronger than those with only a -10 box (Tukey’s honest significant difference [HSD] (Tukey 1949), p=0.039) (**Fig 2E**). Possibly due to low sample sizes, we did not find a significant correlation between promoter strength and the other architectures (Tukey’s HSD).

We next asked whether differences in spacer length also relates to predicted promoter strength. To this end, we identified 128/206 (∼62%) promoters that contain mapped -10 and -35 boxes spaced 16, 17, or 18 bp apart. We first observed that there are approximately twice as many promoters with 17 bp spacers (N=67, ∼52% actual vs 33% expected) than promoters with 16 bp spacers (N=33, ∼26% actual vs 33% expected) or 18 bp-spaced promoters (N=28, ∼22% actual vs 33% expected) (chi-squared test, 2 degrees of freedom, p=2.61×10^-5^). We then compared the strengths of these promoters, but did not find a significant difference between strength and spacer length (ANOVA, f=1.57, p=0.212) (**Fig 2F**). One likely reason for this observation is that promoters with 16 and 18 bp spacers have -10 and -35 boxes that better match their respective consensus sequences than promoters with a 17 bp spacer, i.e. they have higher PWM scores (ANOVA, f=11.2, p=3.49×10^-5^) (**Fig 2G**). To the best of our knowledge, we are not aware of this “compensation” being described elsewhere in the literature.

To summarize (**Fig 2**), almost half (∼46%) of the 206 promoters have just one -10 box and one -35 box, while ∼19% have only a -10 box. Promoters with a -10 and -35 box are stronger than those with only a -10 box. Despite an enrichment of promoters with 17 bp spacers, promoters with spacers of 16, 17, and 18 bp exhibit indistinguishable promoter strengths.

### Spacer length influences mutational robustness

We showed earlier that promoter strength strongly correlates with mutational robustness (see **Fig 1E**), with a coefficient of determination (r^2^) of ∼91%. This suggests that the remaining ∼9% of the variation in robustness is independent of promoter strength. To understand how this residual mutational robustness may be genetically encoded, we normalized the proportion values of each promoter to the trendline shown in Fig 1E to calculate a *normalized robustness* (**Fig 3A**, see **Methods**). By design, normalized robustness does not correlate with promoter strength (Pearson correlation, r=-1.79×10^-15^, p=1.00).

**Figure 3.**
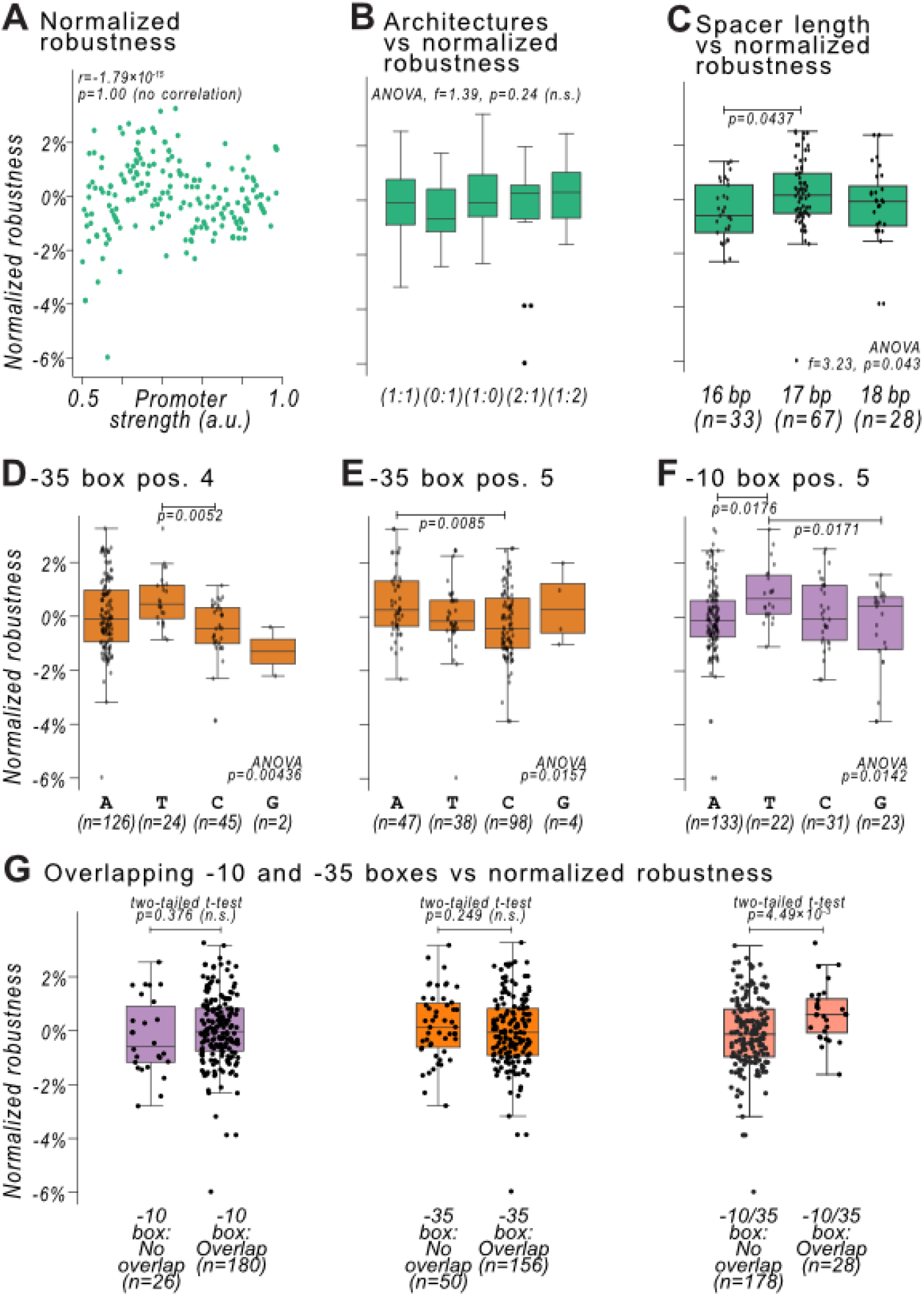
Differences in promoter architecture influence mutational robustness independent of promoter strength. **(A)** Normalized robustness (x-axis) is the mutational robustness value from Fig 1E normalized using equation (1) (Methods). We compared normalized robustness vs predicted promoter strengths for 206 promoters, and tested the null hypothesis that there is no correlation between them (Pearson correlation, r=-1.79×10^-15^, p=1.00). **(B)** We used an analysis of variance (ANOVA) to test whether the normalized robustness scores between different categories of promoters are identical, using only promoter categories with at least 10 promoters. For each category (x-axis), the left and right numbers underneath the box plot refers to the number of -35 boxes and -10 boxes in the promoter (ANOVA, f=1.39, p=0.24, n.s.= not significant). **(C)** For promoters with a -10 and a -35 box spaced 16-18 bp apart (n=128), we used an ANOVA to test the null hypothesis that the mean normalized robustness scores for promoters with different spacer lengths are indistinguishable (ANOVA, f=3.26, p=0.043). P-values above box plots from Tukey’s HSD (Tukey 1949). **(D)** Distributions of normalized robustness scores of promoters with a mapped -35 box. Each column corresponds to the nucleotide identity (A, T, C, or G) at position (pos.) 4 of the -35 box. We used an ANOVA to test the null hypothesis that there is no significant difference between normalized robustness and nucleotide identity at the given position. P-values above box plots from Tukey’s HSD. **(E)** Analogous to (D) but for position 5 of the -35 box. **(F)** Analogous to (D) but for position 5 of the -10 box. **(G)** Normalized robustness promoters with and without overlapping mapped -10 boxes (purple box plots; two-tailed t-test, p=0.376, n.s. = not significant), with and without overlapping mapped -35 boxes (orange box plots; two-tailed t-test, p=0.249, n.s.), as well as with and without overlapping mapped -10 and -35 boxes (salmon-colored box plots; two-tailed t-test, p=4.49×10^-3^).

We then asked whether there are significant differences in normalized robustness for the number of mapped boxes (see Fig 2D), but did not find a significant difference between any pair of groups (ANOVA, f=1.39, p=0.24) (**Fig 3B**). This finding suggests that the number of -10 and -35 boxes does not directly influence mutational robustness, apart from doing so by increasing predicted promoter strength.

Even though the spacing of -10 and -35 boxes (16-18 bp) does not affect promoter strength significantly (see Fig 2F), it may still affect mutational robustness. To find out, we studied the normalized robustness of promoters with mapped -10 and -35 boxes spaced 16, 17, and 18 bp apart, and indeed found a significant difference in robustness (ANOVA, f=3.23, p=0.043) (**Fig 3C**). Specifically, promoters with a 17 bp spacer are more robust than those with a 16 bp spacer (Tukey’s HSD, p=0.0437).

### Individual positions in -10 and -35 boxes are associated with varying robustness

We next asked whether the nucleotide identity at each position in the mapped -10 and -35 boxes influences normalized robustness. To find out, we identified the nucleotide of each validated -10 and -35 box at each of the 6 positions in each box. For any one such nucleotide, we then examined how strongly normalized robustness differs for each of the three other possible nucleotides (**Fig 3D**). For mapped -35 boxes (consensus: TTGACA), we found that promoters with a T in position 4 are more robust than promoters with a C in that position (ANOVA, p=4.36×10^-2^; honest significant difference [HSD] (Tukey 1949), p=5.20×10^-3^), and promoters with an A in position 5 are more robust than promoters with a C in that position (ANOVA, p=1.57×10^-2^; HSD, p=8.50×10^-3^) (**Fig 3E**).

For mapped -10 boxes (consensus: TATAAT), we found that promoters with a T in position 5 are more robust than promoters with an A (ANOVA, p=0.0142; HSD, p=0.0176) or G (HSD, p=0.0171) (**Fig 3F**).

To ensure that these findings were not confounded by differences in promoter strength (which correlates with robustness; see Fig. 1), we tested whether promoter strength differed between the corresponding nucleotide variants using two-tailed *t*-tests. After adjusting the p-values for multiple hypothesis testing (Benjamini Hochberg), we did not find a significant difference in the promoter strengths (**Fig S5**). Ultimately, we demonstrate that specific nucleotides in -10 and -35 boxes can influence mutational robustness independent of predicted promoter strength.

### Promoters with overlapping -10 and -35 boxes are more mutationally robust

Mapped -10 and -35 boxes frequently overlap with each other in promoters (e.g., Fig 1D). When a -10 box overlaps with another -10 box, they can share the same upstream -35 box. Similarly, when a -35 box overlaps with another -35 box, they can share the same downstream -10 box. However, when a -10 box overlaps with a -35 box, this can conversely be two distinct RNA polymerase binding sites. Intuitively, any of these scenarios could potentially influence promoter mutational robustness.

To find out whether the apparent redundancy of overlapping boxes affects mutational robustness, we first categorized promoters into those with and without overlapping -10 boxes, and compared their normalized robustness. Normalized robustness does not differ significantly between the two groups (two-tailed t-test, t=-0.89, p=0.376) (**Fig 3G**). Similarly, it does not differ between promoters with and without overlapping -35 boxes (two-tailed t-test, t=1.16, p=0.249). However, when we looked at promoters with and without overlapping -10 and -35 boxes, we found a significant difference (two-tailed t-test, t=-2.87, p=4.49×10^-3^). Adjusting the p-values for multiple hypothesis testing (Benjamini Hochberg) does not change this significance (q=1.35×10^-2^, FDR=0.05). In sum, overlapping -10 and -35 boxes increases mutational robustness. The most likely explanation for these results is that such promoters comprise two distinct σ70 binding sites, while those with overlapping -10/10 or -35/35 boxes comprise only single σ70 binding sites.

Because overlapping boxes can increase mutational robustness (see Fig 3E), we asked which specific pairs of box instances (see Fig 1A,B) could overlap by 1 bp. From our list of -10 and -35 box instances, we computationally added an A, T, C, or G downstream of each instance, and checked whether this addition created another -10 or -35 box instance. If the addition created a new instance, we call the two instances an *overlapping pair*. We illustrate the different overlapping pairs in **Fig 4A**. We identified overlapping pairs for -35/35 boxes (n=11 pairs), and - 35/10 boxes (n=5 pairs), but none for -10/35 boxes and -10/10 boxes. See **Fig S6** for pairs of instances offset by 2 bp.

**Figure 4.**
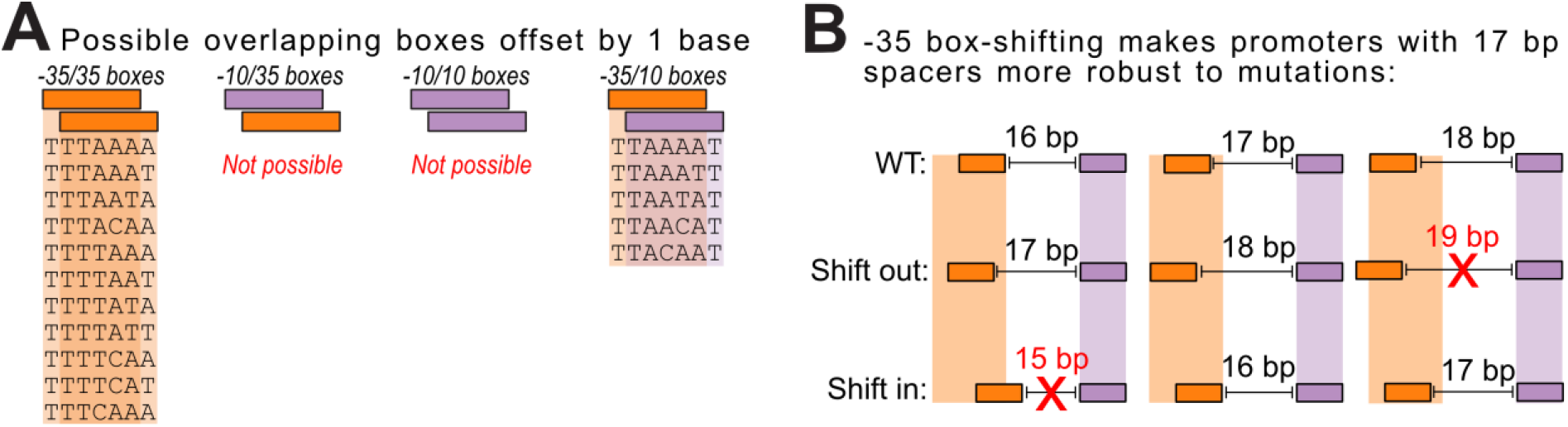
Point mutations can readily shift -35 boxes +/-1 bp upstream and downstream. **(A)** Each column represents one of four scenarios where -10 boxes (purple rectangles) or -35 boxes (orange rectangles) can overlap when offset by one base pair (bp). From left to right: a -35 box overlaps a -35 box, a -10 box overlaps a -35 box, a -10 box overlaps a -10 box, and a -35 box overlaps a -10 box. The DNA sequences in each column correspond to instances of the -10 and -35 boxes that match the respective column. “Not possible” indicates that no combination of -10 and or -35 boxes exists to create the overlap. See **Fig S6** for overlapping boxes offset by 2 bps. **(B)** A schematic illustrating how a -35 box can shift upstream by -1 bp or downstream by +1 bp (along the vertical dimension), and how this shifting can make promoters with a 17 bp spacer more robust than those with 16 or 18 bp. WT: wild-type, starting position. Orange rectangles: -35 boxes, magenta rectangles: -10 boxes. Horizontal lines with flanking vertical bars indicate the spacer distance between the -10 and -35 boxes. Red “X” marks are for spacers at 15 and 19 bp, which do not fit the consensus spacer length.

These observations about overlapping pairs of -35/35 boxes may explain why promoters with 17 bp spacers are more robust (see Fig 3C). With a 17 bp-spaced promoter, mutations upstream or downstream of the -35 box can “shift” the -35 box upstream or downstream and still create a canonical promoter with a spacer of 16-18 bases (**Fig 4B**). Conversely, for promoters with a 16 bp spacer, shifting the -35 box inward is expected to disrupt the promoter, because it creates a 15 bp spacer. The same holds for promoters with an 18 bp spacer, where shifting the -35 box outward creates a 19 bp spacer.

## Discussion

We describe a strong, positive relationship between the strength and mutational robustness of prokaryotic promoters (see Fig 1). From an evolutionary perspective, selection is thought to influence gene expression levels by directly tuning promoter strength. For example, promoters of essential genes drive significantly higher levels of expression than those of nonessential genes (Tsuru et al. 2024). Also, promoters of essential genes are more highly conserved across *E. coli* strains (Lamoureux et al. 2024). Because of the intrinsic relationship between promoter strength and robustness, the traditional viewpoint that selection acts upon promoter strength would suggest that robustness is a product of indirect selection.

Selection, however, may also act directly upon the mutational robustness of a DNA sequence (Forster et al. 2006). Population genetics predicts that this occurs when N_e_×µ×L>>1, where N_e_ is the effective population size, µ is the per nucleotide mutation rate, and L is the length of the mutational target (van Nimwegen et al. 1999; Wagner 2005; Lynch 2011). The -35 and -10 boxes together comprise L=12 base pairs, *E. coli* has ∼N_e_=2×10^8^ individuals (Lynch et al. 2016), and a mutation rate of ∼µ=9×10^-9^ substitutions / nucleotide / generation (Lynch et al. 2016). Thus, N_e_×µ×L=21.6>>1 suggesting direct selection for robustness is at least possible (Wagner 2005).

Because selection can, in theory, act directly on the mutational robustness of a promoter, we cannot rule out the counterintuitive possibility that some strong promoters are actually selected for their *robustness* and not necessarily their strength. This alternative explanation would also explain why promoters for essential genes drive high expression (Tsuru et al. 2024). Furthermore, it may also help to explain why mRNA levels correlate only moderately with protein levels in *E. coli* (Taniguchi et al. 2010; Riba et al. 2019; Peterman et al. 2025). Specifically, protein levels are largely shaped by post-transcriptional regulation such as codon usage (Han et al. 2010; Liu et al. 2018), RNA secondary structures (Mandal and Breaker 2004), and ribosome binding site affinity (Goodman et al. 2013; Kosuri et al. 2013; Cambray et al. 2018). Possibly the key selective pressure on protein expression is not just the fine-tuning of mRNA levels, but also the consistent and robust production of transcripts.

One additional observation also supports the possibility of selection for robustness. Nearly twice as many promoters have a 17 bp spacer than a 16 bp or 18 bp spacer, despite indistinguishable promoter strengths (see Fig 2F). The overrepresentation of 17 bp spacers therefore cannot be explained by promoter strength alone. Instead, it may reflect the fact that 17 bp promoters are more robust to mutations (see Fig 3C and Fig 4B). We urge caution in this interpretation, as it could also be a bias of the thermodynamic model to favor promoters with 17 bp spacers.

Regardless of possible selective pressures, we found that promoters differ in the number of -10 and -35 boxes, which can contribute to their activity (see Fig 2D,E). This observation is consistent with previous studies which estimate that ∼20% of all promoters in the *E. coli* genome do not have a -35 box (Mitchell et al. 2003), and that genomic promoters can encode multiple -10 and - 35 boxes (Huerta and Collado-Vides 2003). While we initially hypothesized that these extra boxes contribute to robustness, our analysis does not support this (see Fig 3B). The extra boxes may exist either because they are remnants of previously active but now eroded promoters (Huerta and Collado-Vides 2003; Bradley et al. 2010; Krieger et al. 2022), or by chance alone. The latter possibility is not far-fetched, given that synthetic random DNA sequences often harbor -10 and - 35 boxes. Some 10% of synthetic DNA even encodes a functional promoter (Yona et al. 2018; Lagator et al. 2022).

Altogether, we demonstrate that there is an intrinsic relationship between promoter strength and mutational robustness. Because of this relationship, we cannot rule out the possibility that selection acts upon both promoter strength as well as mutational robustness. Mutational robustness can be further increased, independent of promoter strength, by equipping promoters with particular instances of the -10 and -35 boxes, encoding overlapping -10 and -35 boxes, and having -10 and -35 boxes spaced exactly 17 bp apart. Our analyses provide insights into the evolutionary underpinnings of promoter strength and robustness, as well as possible strategies for designing more robust promoters in synthetic biology.

## Data availability

We provide a Jupyter notebook with the python scripts to recreate the figures and analyses in this document at a local Github repository: https://github.com/tfuqua95/robust_promoters

**Data S1 (separate file)**. A csv file with the predicted promoter strengths for the 206 promoters and their DNA sequences.

**Data S2 (separate file)**. A csv file with the predicted promoter strengths for all single point mutant sequences.

**Data S3 (separate file)**. A csv file with the coordinates of the mapped -10 and -35 boxes, original p-values, their corresponding corrected q-values, and the contribution each box has on the promoter strength. This information is also compiled into Data S1 for each promoter sequence.

## Materials and methods

### Position weight matrices (PWMs)

We obtained PWMs for -10 and -35 boxes using a list of -10 and -35 boxes from (Belliveau et al. 2018), which were originally derived from (Tierrafría et al. 2022). They are derived specifically from the -10 and -35 boxes of promoters that were either experimentally confirmed or from promoters with “strong” computational and experimental evidence. See https://regulondb.ccg.unam.mx/manual/help/evidenceclassification for more information behind these classification terms or ref (Weiss et al. 2013).

We converted these sequences into PWMs using the Biopython.motifs package (Cock et al. 2009). We assumed a background distribution of 25% for each nucleotide (A, T, C, and G). PWMs assign scores, measured in bits or nats, to query sequences. A higher score indicates a greater likelihood that the sequence interacts with a specific DNA binding protein. Because PWM scores can vary widely, determining a universal threshold for defining a functional transcription factor binding site is not always straightforward. We used the widely used (Doniger et al. 2005; Dresch et al. 2025; Lee et al. 2025) “Patser threshold,” which corresponds to the motif’s total information content in nats (Hertz and Stormo 1999). Specifically, our -35 box PWM has an information content of 3.39 nats, while the -10 box has 3.98 nats. We drew the sequence logos in Fig 1A-B using Logomaker (Tareen and Kinney 2020).

### Statistical thermodynamic model and the 206 genomic promoters

We acquired 1,997 sigma70 promoter DNA sequence from Regulon DB (Tierrafría et al. 2022), of which 55 (∼3%) are considered to be “confirmed”, 785 (∼39%) have “strong” evidence of being promoters, and 1’161 (∼58%) have “weak” evidence of being promoters. See https://regulondb.ccg.unam.mx/manual/help/evidenceclassification for more information behind these classification terms or ref (Weiss et al. 2013).

For each 81 bp promoter sequence, we computationally concatenated 22 Gs upstream and 22 Gs downstream to match the thermodynamic model’s preference length for a 125 bp query sequence (Lagator et al. 2022). For each sequence, we then used the model (standard thermodynamic model with flexible spacers and cumulative binding effects: stm+flex+cumul) to calculate the probability of the sequence being a promoter, and identified 206 promoters with a score greater than 0.5 a.u.

Of the 206 promoters, 6 (∼3%) are experimentally “confirmed”, 96 (∼47%) have a “strong” prediction confidence, and 104 (∼50%) have a “weak” prediction confidence. There are significantly more “strong” predictions in the set of 206 promoters compared to the full set of 1,997 promoters (Fischer’s exact test, odds = 0.74, p=0.044), and significantly fewer “weak” predictions compared to the full set of 1,997 promoters (Fischer’s exact test, odds=1.36, p=0.039).

See **Data S1** for the promoter sequences we analyzed and their predicted promoter scores. We applied the same procedure to the saturation mutagenesis libraries, whose scores and sequences can be found in **Data S2**.

### Mapping -10 and -35 boxes

We identified -10 and -35 boxes in the wild-type 206 promoter sequences using position-weight matrices (PWMs) as described in the subsection *Position Weight Matrices (PWM)*. For each of these boxes, we computationally partitioned the mutant sequences into two groups, those with a mapped box, and those without a mapped box. We then tested the null hypothesis that the respective promoter strengths from the thermodynamic model (see *Statistical thermodynamic model and the 206 genomic promoters*) were the same using a Mann-Whitney U (MWU) test. We performed this test only if each of the two groups contained at least 5 mutant sequences. We used the Benjamini-Hochberg procedure to account for multiple-hypothesis testing (Benjamini and Hochberg 1995). Because the MWU test is sensitive to small differences in distributions, we only kept significant matches that exhibited at least a median difference of 0.1 a.u. between both groups for further analyses. See **Data S3** for all of the mapped -10 and -35 boxes. The coordinates of each -10 and -35 box for each promoter are also available in **Data S1**.

### Normalized robustness scores

We calculated a line of best fit between P_pro_ and promoter strengths in Fig 1E using `scipy.stats.linregress` (Virtanen et al. 2020), fitting the linear equation to equation (1):

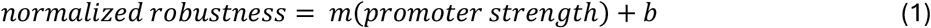

which resulted in fitted coefficients of m=0.304 and b=0.65. We used equation (1) to calculate the normalized robustness scores for the 206 promoters based on their promoter strengths (see *Lagator model and the 206 genomic promoters*).

## Acknowledgements

We thank all members of the Wagner group for discussions. We also thank Baxter for his own robustness.

---

**Figure S1.**
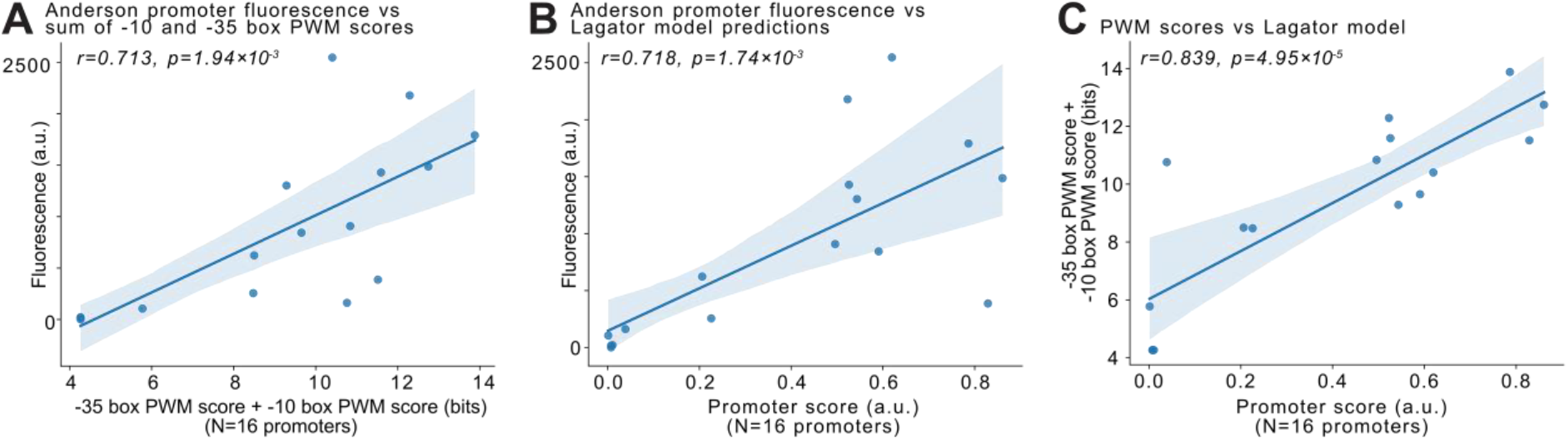
Promoter model predictions significantly correlate with experimental results. **(A-C)** The Anderson promoter collection is an iGEM promoter library consisting of 19 promoter sequences (J23119 and J23110-J23118), of which the activity of 16 sequences has been measured using a red fluorescent protein (RFP) reporter plasmid in *E. coli* (https://parts.igem.org/Part:BBa_J23110). **(A)** RFP fluorescence (arbitrary units, a.u.) vs the sum of the -35 box and -10 box position weight matrix (PWM) scores in information theoretical units (bits) (Pearson correlation, r=0.713, p=1.94×10^-3^). **(B)** RFP fluorescence (a.u.) vs predicted promoter strength using the Lagator Model (see Methods; Pearson correlation, r=0.718, p=1.74×10^-3^). **(C)** The sum of the -35 box and -10 box PWM scores vs promoter strength (Pearson correlation, r=0.839, p=4.95×10^-5^). This figure shows that PWM scores and the thermodynamic model can predict promoter activity, and that the predictions are comparable to each other.

**Figure S2.**
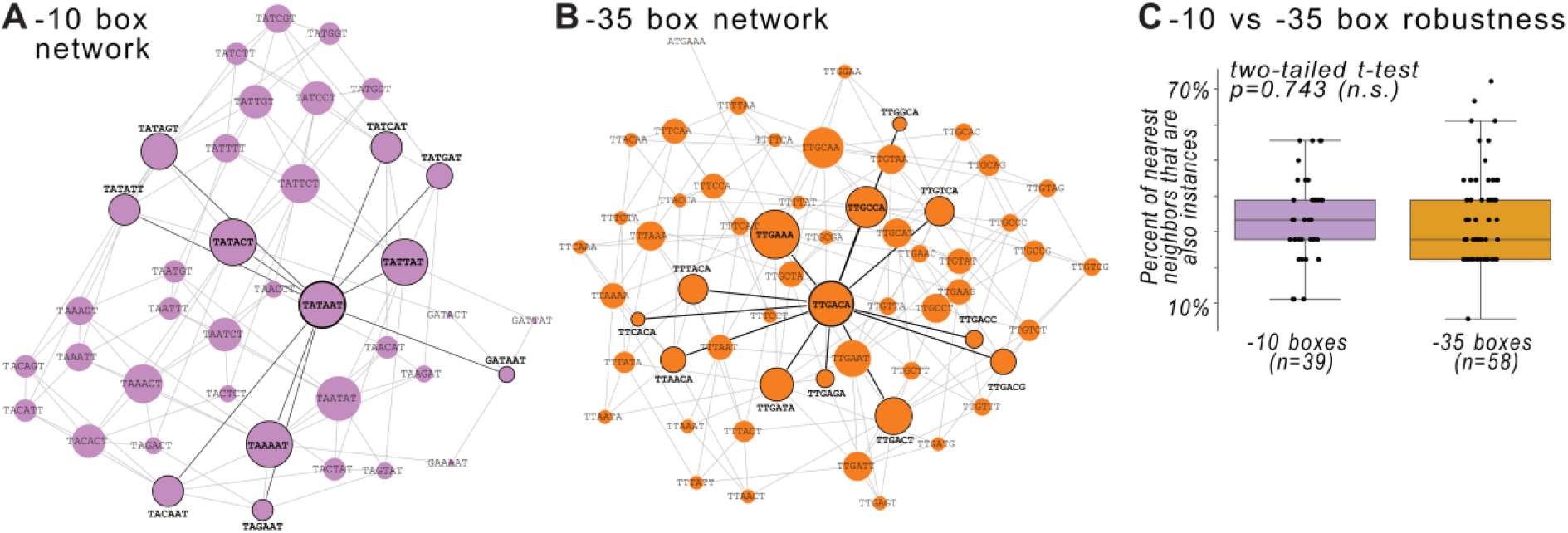
-10 and -35 box instances. **(A)** The genotype network for 39 position-weight matrix (PWM) predicted instances of the -10 box (see Fig 1). Each circle (node) represents a unique genotype. Each straight line (edge) connects two boxes that differ in a single nucleotide. The central node outlined in black is the canonical consensus sequence. Networks visualized using Gephi (Bastian et al. 2009). **(B)** Analogous to (A) but for 59 instances of the - 35 box. **(C)** Box plots comparing the percentage of neighbors for each box instance that are also instances (left: -10 box instances, n=39; right: -35 box instances, n=59). We test the null hypothesis that there is no significant difference between the distributions with a two-tailed t-test (p=0.743, n.s. = not significant).

**Figure S3.**
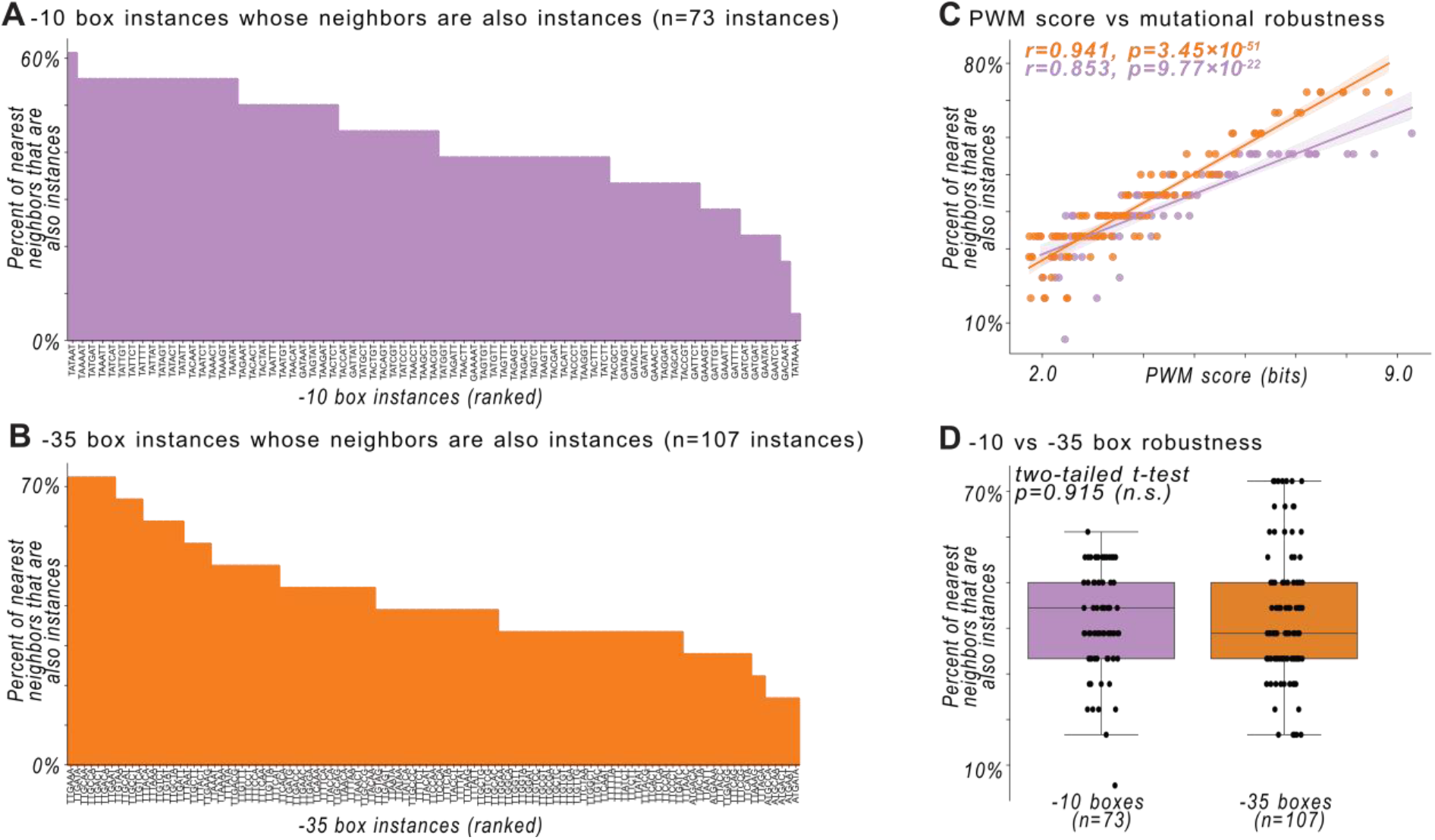
Robustness with a lower PWM threshold. **(A)** We calculated all possible DNA sequence instances that the PWM classifies as a -10 box using half of the Patser threshold (-10 box ≥ 1.990 nats). For each instance of such a box, we plot the percentage of its nearest mutational neighbors (out of 6×3=18 possible neighbors) that are also -10 box instances. **(B)** Analogous to (A) except for the -35 box using half of the Patser threshold (-35 box ≥ 1.695 nats). **(C)** The PWM scores in bits for each box instance vs the percentage of a box’s nearest neighbors that are also an instance when the Patser threshold is halved. Orange circles: -35 box instances, magenta circles: -10 box instances. We tested the null hypothesis that there is no correlation between the percentages and PWM scores using a Pearson correlation (-35 box: r=0.941, p=3.45×10^-51^; -10 box: r=0.853, p=9.77×10^-22^). **(D)** Box plots comparing the number of neighbors for each box instance (left: -10 box instances, n=73; right: -35 box instances, n=107). We test the null hypothesis that there is no significant difference between the distributions with a two-tailed t-test (p=0.915, n.s. = not significant).

**Figure S4.**
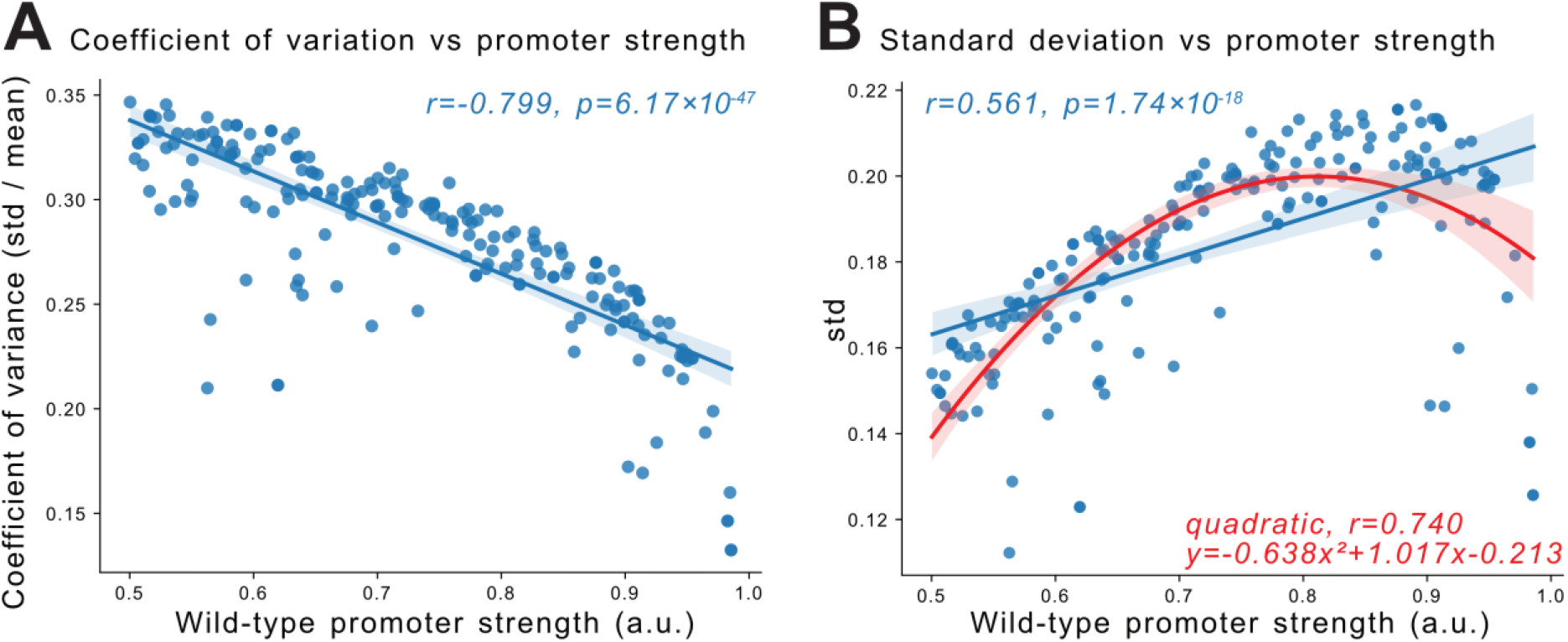
Alternative definitions of robustness. **(A)** A scatterplot capturing the relationship between promoter strength as predicted using the thermodynamic model for 207 promoter sequences (x-axis) vs the coefficient of variance for the mutant sequences (y-axis). We test the null hypothesis that there is no correlation between the variables using a Pearson correlation (r=-0.799, p=6.17×10^-47^). **(B)** Analogous to (A) but for promoter strength vs the standard deviation for the mutant sequences(y-axis) (Pearson correlation, r=0.561, p=1.74×10^-18^). We also fit the relationship to a quadratic equation (y=-0.638x^2^-1.017x-0.213), with a coefficient of determination (r) = 0.740, suggesting the quadratic relationship better explains the relationship than the linear one does.

**Figure S5.**
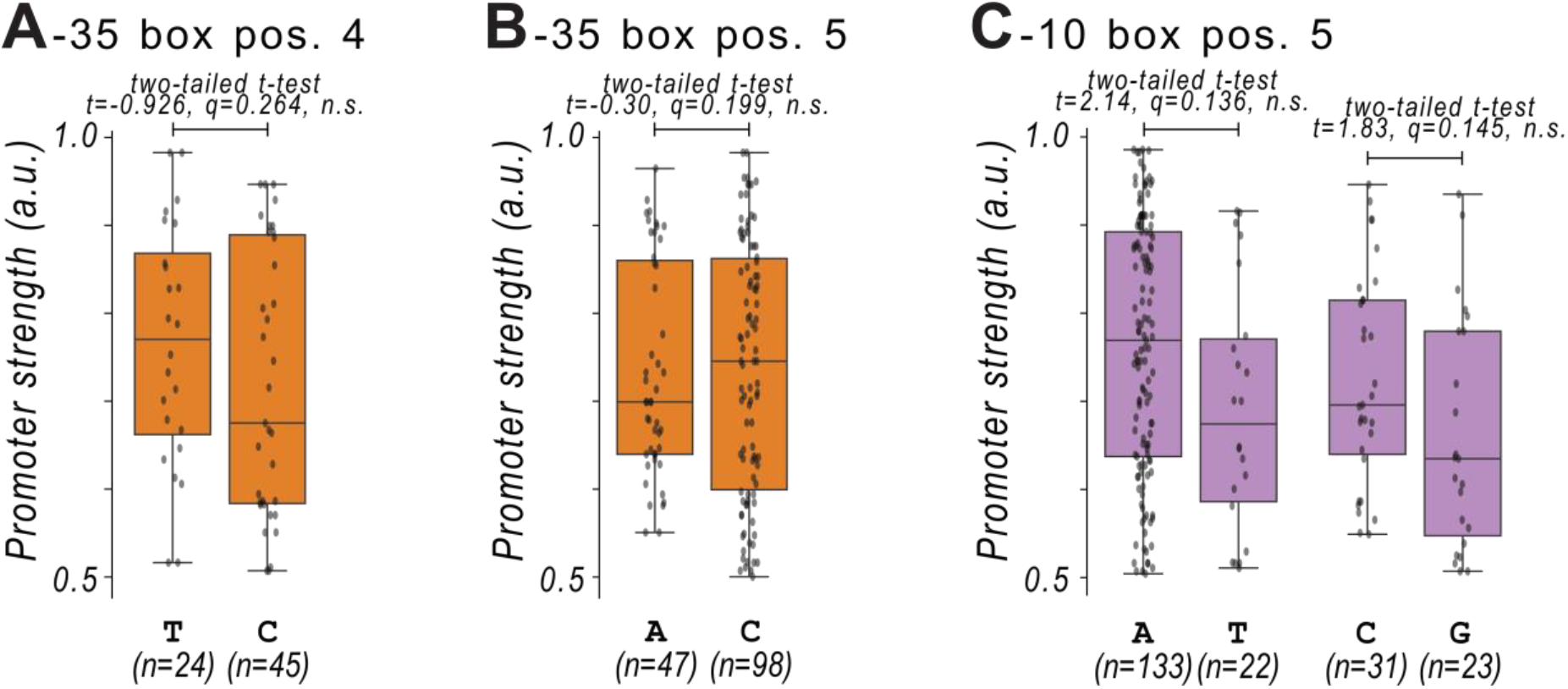
Promoter strength vs robust positions. **(A)** Distributions of promoter strengths (arbitrary units, a.u.) with a mapped -35 box. Each column corresponds to the nucleotide identity (A, T, C, or G) at position (pos.) 4 of the -35 box that we identified in Figure 3 to be significantly more robust. We test the null hypothesis that there is no significant difference in the promoter strengths using a two-tailed t-test, and calculate corresponding q-values to account for multiple hypothesis testing using the Benjamini-Hochberg procedure (two-tailed t-test, t=-0.926, p=0.357, q=0.264, n.s.=not significant). **(B)** Analogous to (A) but for position 5 of the -35 box (two-tailed t-test, t=-0.30, p=0.976, q=0.199). **(C)** Analogous to (A) but for position 5 of the -10 box (two-tailed t-test, A vs T: t=2.14, p=0.034, q=0.136; C vs G: t=1.83, p=0.073, q=0.145).

**Figure S6.**
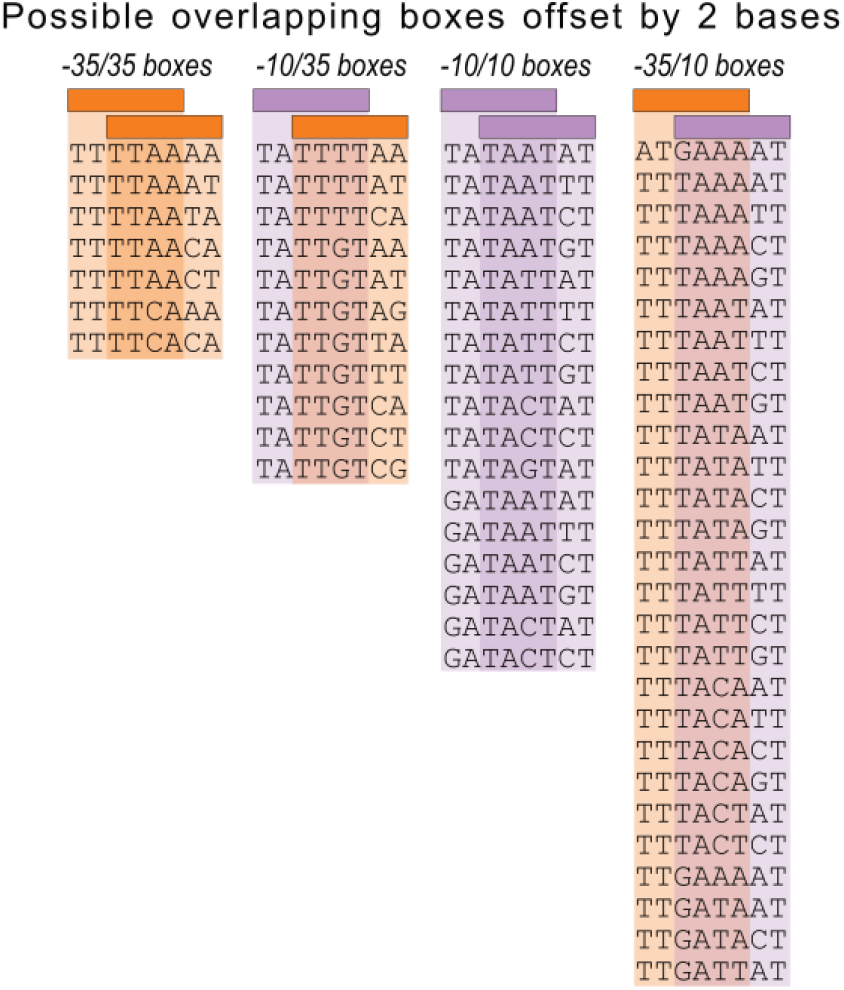
Possible overlapping boxes offset by 2 bases. Each of the four columns represents one of four possible scenarios where -10 boxes (purple rectangles) and -35 boxes (orange rectangles) can overlap when offset by one base pair (bp) (from left to right): a -35 box overlaps a -35 box, a -10 box overlaps a -35 box, a -10 box overlaps a -10 box, and a -35 box overlaps a -10 box. The DNA sequences in each column correspond to instances of the -10 and - 35 boxes that match the respective scenario. “Not possible” indicates that no combination of -10 and or -35 boxes exists to create the overlap. See **Fig 4A** for overlapping boxes offset by 1 bp.

